# Cross-feeding interactions between *Fusobacterium nucleatum* and the glycan forager *Segatella oris*

**DOI:** 10.1101/2025.06.18.660387

**Authors:** Joshua R. Fletcher, Ryan C. Hunter

**Affiliations:** Department of Population Health and Pathobiology, North Carolina State University College of Veterinary Medicine, Raleigh, NC 27606; Department of Microbiology & Immunology, Jacobs School of Medicine and Biomedical Sciences, University at Buffalo, Buffalo, NY 14203

**Keywords:** *Fusobacterium nucleatum*, *Segatella oris*, *Prevotella oris*, cross-feeding, glycans, microbiome, chronic airway disease, transcriptomics

## Abstract

*Fusobacterium nucleatum* is a common member of the oral microbiota frequently associated with extra-oral infections and diverse polymicrobial environments, including chronic airway diseases and colorectal tumors. Yet, its interactions with co-colonizing microbiota remain poorly defined. Here, we investigate cross-feeding interspecies dynamics between *F. nucleatum* and *Segatella oris,* a glycan-foraging anaerobe enriched in airways and gastrointestinal tumors. Using broth cultures, cell-free supernatants, and co-culture on primary human airway epithelial cells, we identify bidirectional interactions that shape nutrient acquisition, biofilm formation, gene expression, and host responses. While mucin or *S. oris* supernatants modestly enhanced *F. nucleatum* growth, both conditions triggered transcriptional remodeling, including induction of the *nan* operon for sialic acid catabolism, suggesting reliance on glycan degradation by *S. oris.* Conversely, *S. oris* exhibited differential expression of multiple polysaccharide utilization loci (PULs) when exposed to *F. nucleatum* or its metabolites. Biofilm formation by *F. nucleatum* was strongly inhibited by *S. oris,* indicative of antagonistic interactions. Dual and triple RNA-seq revealed that epithelial responses were predominately shaped by *F. nucleatum,* with enrichment of inflammatory and cancer-associated pathways; however, co-colonization with *S. oris* modulated the magnitude and specificity of host gene expression. These findings demonstrate that glycan-mediated cross-feeding and microbial interactions shape the physiology and pathogenic potential of *F. nucleatum* in mucosal environments. This work underscores the importance of modeling polymicrobial communities under host-relevant conditions to better understand pathobiont behavior at the epithelial interface.

**Importance:** *Fusobacterium nucleatum* is increasingly recognized as a pathobiont in mucosal diseases, including colorectal cancers and chronic airway infections, yet its functional interactions with co-colonizing microbiota remain poorly understood. Here, we demonstrate that *F. nucleatum* engages in bidirectional interactions with *Segatella oris,* a glycan-foraging anaerobe also enriched in mucin-rich environments. Through nutrient cross-feeding, antagonism, and transcriptional modulation, these interactions shape bacterial behavior and the host epithelial response. Notably, glycan degradation by *S. oris* enables *F. nucleatum* access to sialic acids, while *F. nucleatum* suppresses expression of multiple polysaccharide utilization loci in *S. oris,* revealing a reciprocal ecological influence. Co-colonization of the airway epithelial surface also modulates gene expression linked to inflammation and cancer. These findings advance our understanding of polymicrobial dynamics at mucosal interfaces and highlight the importance of incorporating microbe-microbe-host interactions into reductionist models of infection and disease.

## Introduction

*Fusobacterium nucleatum* is a Gram-negative member of the oral microbiota that contributes to biofilm assembly, architecture, and community organization (1, 2). Although best known for its roles in dental plaque and periodontal disease, *F. nucleatum* is implicated in a range of extraoral infections and is frequently detected in tumors across multiple cancer types (3–6). While it can function as both a commensal and pathogen, *F. nucleatum* is often found in diverse polymicrobial host environments, including colorectal tumors and the airways of individuals with chronic rhinosinusitis (CRS), cystic fibrosis (CF), or chronic obstructive pulmonary disease (COPD) (7–14). This raises questions about whether and how *F. nucleatum* interacts with co-colonizing microbiota and how these interactions shape its behavior and pathogenic potential. Equally important is understanding how *F. nucleatum* influences the activity of other taxa in these communities, many of which also exhibit context-dependent commensal or pathogenic behavior (15).

*Segatella oris* (formerly *Prevotella oris*) is also a member of the oral microbiota, though less is known about its physiology or microbial interactions. Notably, *S. oris* supernatants inhibit growth of the airway pathogen *Moraxella catarrhalis in vitro*, suggesting a connection between *S. oris* residence in the upper airways and colonization resistance against some pathogens (16). However, *S. oris* also exhibits pathogenic traits, including hemolysin production with activity against human erythrocytes (17). Like *F. nucleatum*, it is detected in extraoral infections, gastric tumors, CRS and CF airway mucus, and is co-enriched with *F. nucleatum* in the oral microbiomes of cancer patients with chemotherapy-induced oral mucositis (9, 18, 19). More broadly, both *Fusobacterium* and *Prevotella* genera are co-detected in esophageal tumors (20), suggesting that cross-feeding interactions between these taxa may facilitate colonization and promote disease.

Although environmental conditions vary across infection sites, shared features such as host glycans, proteoglycans, and glycosaminoglycans are common. This is particularly true in mucus-laden airways of patients with chronic respiratory diseases and the mucus layer lining the intestinal tract, where *F. nucleatum* has been detected (21). *F. nucleatum* primarily ferments amino acids to generate ATP, although glucose and fructose metabolism have also been reported (22–26). While it produces intracellular polyglucose granules *in vitro*, the genetic basis and functional significance of this pathway, including possible links to virulence, remain unclear (27). Notably, *F. nucleatum* lacks sialidases and polysaccharide utilization loci (PULs), but can scavenge sialic acids liberated by glycan foraging microbes (28), potentially enhancing community persistence through cross-feeding. In contrast, the *S. oris* NCTC 13071 genome encodes 27 predicted PULs, likely involved in sensing, importing, and degrading of complex polysaccharides such as mucins and other host glycans (29, 30).

Given their frequent co-occurrence in mucin-rich environments *in vivo* and the possible complementarity of their metabolic repertoires, we hypothesized that *F. nucleatum* and *S. oris* engage in cross-feeding interactions. To test this, we examined how each species adjusts its growth and gene expression to mucin glycoproteins and each other’s metabolic byproducts. We found that while neither mucins nor *S. oris* supernatant substantially altered the growth of *F. nucleatum*, both conditions triggered significant transcriptional changes. Conversely, *S. oris* exhibited distinct growth dynamics and widespread transcriptional responses in mucin and *F. nucleatum* supernatants. Although *F. nucleatum* forms robust biofilms, biofilm formation was greatly reduced in *S. oris* supernatants and in co-culture, despite biofilm growth in mucin or control media. We further extended these analyses using co-culture of each species, individually and together, on the apical surface of primary human airway epithelial cells to model aspiration-based colonization relevant to CF, CRS, and COPD. Dual (host-*S. oris,* host-*F. nucleatum*), and triple (host-*S.* o*ris-F. nucleatum)* RNA sequencing revealed that host-associated growth conditions strongly influence bacterial gene expression, including unexpected variability in *S. oris* PUL expression mediated by *F. nucleatum*. Host transcriptomic responses were primarily driven by *F. nucleatum,* with prominent induction of pro-inflammatory mediators *TNFA* and *TNFAIP2*, as well as several genes linked to cancer. Together, these data capture key information about bacterial interactions on relevant host cell types and highlight how co-colonizing, cross-feeding partners may modify each other’s physiology and influence host responses.

## Materials and methods

### Bacterial strains and media conditions

*Fusobacterium nucleatum* subsp. *nucleatum* strain ATCC 25586 and *Segatella oris* strain NCTC13071 were propagated anaerobically in a Coy anaerobic chamber using BBL *Brucella* broth (BD) or 1.5% agar plates supplemented with hemin (250 μg/mL) and vitamin K (50 μg/mL) (Hardy Diagnostics). A semi-defined control medium was prepared by combining Brucella Broth with minimal salts (50:50) as previously described (31). A minimal mucin medium (MMM) was generated by autoclaving porcine gastric mucin (PGM, Sigma) at 30 g/L in water, diluting to 15 g/L in 2X minimal salts, and combining 1:1 with Brucella broth to yield the final mucin-containing experimental medium (hereafter referred to as mucin medium), adapted from Flynn et al. (32) with dialysis steps omitted. Cell-free supernatants (CFS) were prepared in biological triplicate by culturing each strain for 48 h in mucin medium (10 mL) under anaerobic conditions. Cultures were centrifuged at 4,000 rpm at 4℃ for 20 min, and the resulting supernatants were then passed through 0.22 μm filters. CFS were stored at −80℃ and thawed once immediately prior to use.

### Growth curves

*F. nucleatum* and *S. oris* were grown overnight in Brucella Broth supplemented with hemin (0.35 mg/mL) and vitamin K (0.05 mg/mL), then subcultured 1:5 in fresh medium and grown for an additional 4h. Optical density at 600 nm (OD600) was determined via spectrophotometry and adjusted to 0.01 in the respective growth medium. 200 μL of each culture was added to individual wells in a clear, flat-bottomed 96-well plate in technical triplicate across three biological replicates. Plates were sealed with a Breathe-Easy gas-permeable membrane and incubated in a Tecan Sunrise plate reader in the anaerobic chamber at 37℃ for 72h. OD600 was recorded hourly following 5 seconds of linear shaking.

### Biofilm assays

Biofilm assays were performed in parallel using identical media conditions as described for growth curves, following the protocol of Merritt et al (33). Briefly, microtitre plates were inoculated with each species individually or in co-culture and incubated anaerobically at 37℃ for 48 h. Following incubation, plates were removed from the chamber and planktonic cultures were discarded. Plates were washed three times with 250 μL water, stained with 200 μL 0.1% crystal violet (in water) for 15 minutes, then washed again three times and air-dried overnight. Crystal violet was solubilized with 200 μL of 30% acetic acid, and absorbance at 550 nm was determined using a BioTek Synergy H1 microplate reader. Wells containing sterile media were used as negative controls for background subtraction.

### Colonization of primary human airway epithelial monolayers

Normal human bronchial epithelial (NHBE) cells (Lonza Bioscience), obtained from healthy donors were expanded B-ALI Growth Basal Medium (Lonza) and seeded on 6.5 mm Transwell inserts (0.4 um pore; STEMCELL Technologies). Upon reaching confluency after ∼2-4 days post-seeding, apical medium was removed, and basolateral medium was replaced with B-ALI Differentiation Basal Medium (Lonza). Cells were maintained at air-liquid interface (ALI) for 3-4 weeks at 37℃ and 5% CO_2_ in a humidified incubator.

Overnight bacterial cultures were grown in Brucella Broth supplemented with hemin and vitamin K. These were subcultured 1:5 in fresh medium and grown to OD600 = 0.5. Cultures were diluted 1:10 into infection medium (DMEM supplemented with 2% FBS, 10mM HEPES, 0.1 mM nonessential amino acids, 4 mM L-glutamate, and 1mM sodium pyruvate). 100 μL of bacterial suspension was applied to the apical surface of each Transwell and incubated for 4 h at 37℃ under anaerobic conditions. After incubation, apical supernatants were gently removed by pipette, and co-cultures were maintained for an additional 24h prior to RNA extraction.

### RNA extraction

Each species was grown in 10 mL of control, mucin, and CFS media in 15 mL conical Falcon tubes for 24h and collected by centrifugation at 4,000 rpm for 20 min at 4℃. Pellets were dissolved by gentle pipetting in 1 mL of TRIzol Reagent (ThermoFisher). For airway epithelial co-cultures, 250 μL TRIzol was added to the apical surface and a 1 mL pipette tip was used to scrape cellular material from the transwell surface. Scraping was performed four times per insert to yield 1 mL of total lysate in TRIzol. All TRIzol samples were incubated for 5 minutes at room temperature, followed by addition of 200 μL chloroform, hand agitation for 15 s, followed by another 5-minute incubation on the benchtop. Phase separation was performed by centrifugation at 12,000 rpm for 15 minutes at 4℃. The aqueous phase was mixed 1:1 with 95% ethanol and RNA was column purified with the Zymo Clean & Concentrator-5 kit including an on-column DNase-I treatment according to manufacturer’s instructions.

### RNA sequencing and analysis

Total RNA from broth cultures was submitted to Seq Center (Pittsburgh, PA), where rRNA depletion was performed with the Illumina Stranded Total RNA Prep with Ribo-Zero Plus Microbiome kit prior to library preparation and sequencing (2 x 150 bp). RNA from bacterial and epithelial co-cultures was sent to SeqCoast Genomics (Portsmouth, NH). rRNA depletion was also performed on these samples prior to Illumina sequencing (2 x 150 bp). Sequencing was performed on the Illumina NextSeq2000 platform with a 300 cycle flow cell kit. Read quality was assessed using FastQC (https://www.bioinformatics.babraham.ac.uk/projects/fastqc/). Given the consistently high quality and the risk of biasing downstream analyses (34), no read trimming was performed.

For bacterial RNA seq, coding sequences for all annotated genes and riboswitches, gene names, and locus tags were extracted from the *F. nucleatum* ATCC 25586 (NZ_CP028101) and *S. oris* NCTC 17031 (NZ_LR134384) genomes in Geneious Prime (2024.0.7) and converted to FASTA format via the “Tabular-to-fasta” tool via the Galaxy server (https://usegalaxy.org/), and indexed with Salmon (35) for quasi-mapping of reads. For host RNA-seq, the *Homo sapiens* transcriptome (GRCh38.p14, release 45) was retrieved from Gencode (https://www.gencodegenes.org/human/release_45.html). Transcriptome indices were built in Salmon, to which reads were quasi-mapped. The quant.sf files generated by Salmon were imported into RStudio via the tximport package (36) and differential expression analysis was performed with DESeq2 (37). The threshold for differential expression was a log2 fold change ≥1 at an adjusted p-value <0.05. Code for each analysis is available at https://github.com/Hunter-Lab-UMN/Fletcher_FnSo_2025.

### Data availability

All sequencing files are available at the NCBI Sequence Read Archive (SRA) under accession number PRJNAXXXXX.

## Results

### Growth and biofilm formation of *F. nucleatum* and *S. oris* are influenced by mucin and cross-feeding *in vitro*

*F. nucleatum* and *S. oris* were cultured under three conditions: (i) control medium, (ii) control medium supplemented with porcine gastric mucin (hereafter “mucin medium” or “mucin”), or (iii) cell-free supernatants (CFS) derived from each species grown in mucin for 48h. In control medium, *F. nucleatum* exhibited distinct growth phases with a pronounced exponential phase followed by stationary and death phases, with optical density remaining consistent after ∼36h. Exponential growth rates were similar across conditions; however, cultures grown in mucin or *S. oris* CFS reached higher stationary-phase densities and exhibited a slower decline during death phase compared to controls (∼12-36h, **Fig. 1A**). This elevated density in mucin was sustained until ∼60h, after which it declined to control levels. Area under the curve (AUC) analysis reflected this trend, with mucin medium showing a modest, though not statistically significant, increase relative to the control (p=0.0951, **Fig. 1B).** In contrast, *F. nucleatum* density in *S. oris* CFS declined sharply after 48h, yielding an AUC comparable to the control.

**Fig. 1.**
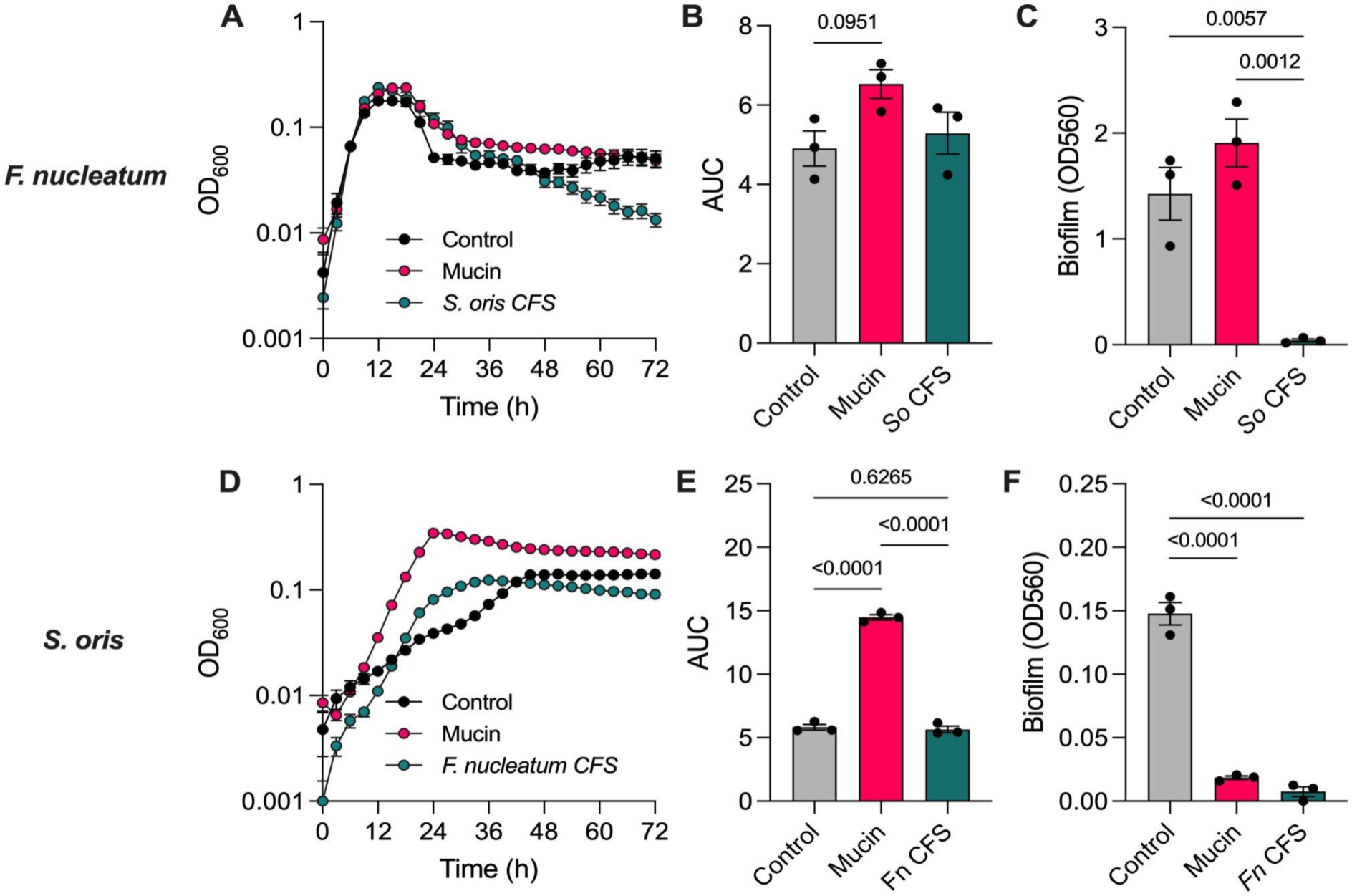
Mucin and interspecies metabolite exchange influence growth and biofilm formation of *F. nucleatum* and *S. oris*. (**A**) Growth curves of *F. nucleatum* in control medium, mucin medium, and *S. oris* cell-free supernatants (CFS) (n=3). (**B**) Area under the curve (AUC) analysis for *F. nucleatum* growth. (**C**) Biofilm formation by *F. nucleatum* in each condition after 48h, quantified by crystal violet staining. (**D**) Growth curves of *S. oris* in control medium, mucin medium, and *F. nucleatum* CFS. (**E**) AUC analysis for *S. oris* growth. (**F**) Biofilm formation by *S. oris* across all conditions. Data represent the mean of three biological replicates +/-standard error of the mean (n=3). Statistical significance in panels B,C,E,F was determined using ordinary one-way ANOVA with Tukey’s multiple comparisons test.

Biofilm formation by *F. nucleatum* was robust in both control and mucin media, with a trend toward increased biofilm in mucin. However, biofilm growth was nearly abolished in *S. oris* CFS (**Fig. 1C**). This suppression is unexpected given *F. nucleatum’s* well-established role as a so-called “bridging” species in the highly structured and spatially stratified polymicrobial biofilms of the oral cavity (38).

In contrast, *S. oris* exhibited markedly different growth dynamics across conditions. Growth in control medium was slow with apparent diauxie following a brief stationary phase at ∼24h, a second exponential phase, and final plateau at ∼40h (**Fig. 1D**). In mucin medium, *S. oris* grew much more rapidly, reaching a higher maximum density and exhibiting a three-fold increase in AUC compared to the control (**Fig. 1E)**. These findings are consistent with genomic predictions of glycan utilization via PULs. Growth in *F. nucleatum* CFS yielded a faster growth rate than control medium but lower stationary phase density than mucin; AUC for CFS and control were comparable. Biofilm formation by *S. oris* was minimal in the control medium and nearly undetectable in mucin or CFS conditions (**Fig. 1F**). In dual-species cultures grown in control or mucin media, biofilms phenocopied the minimal single-species biofilms of *S. oris* (**Suppl. Fig. 1**).

### *F. nucleatum* and *S. oris* transcriptomes are modulated by mucin, cross-feeding, and interactions with human airway epithelia

Both species exhibited growth phenotypes responsive to the nutritional composition of each medium (**Fig. 1)**. Mucin supplementation modestly enhanced *F. nucleatum* growth but had a pronounced effect on *S. oris,* suggesting species-specific nutrient utilization. Growth in CFS derived from either species grown in mucin medium was comparable to or greater than growth in the control medium, indicating that neither species fully exhausted nutrients essential to the other. Interestingly, *S. oris* growth was diminished in *F. nucleatum* CFS compared to mucin, despite *F. nucleatum* lacking known glycan-degrading capabilities. Given their potential clinical relevance and prevalence of each species in chronic airway disease, we also sought to connect broth culture data to more physiologically relevant conditions with host airway epithelia. To model how each species influences the host airway response to colonization, we employed a Dual Oxic-Anoxic Co-Culture (DOAC) model recently developed by our group, in which oxygen-dependent airway epithelial cells are supplied oxygenated blood gas basolaterally while exposing their apical surfaces to the anaerobic chamber environment (39). This advancement allows for colonization and growth of obligate anaerobes on epithelial cells, enabling study of host-pathogen-microbiota interactions. We performed RNA-seq on each bacterium in control, mucin, and CFS media, as well as dual- and triple-RNA-seq on primary normal human bronchial epithelial (NHBE) cells challenged with each bacterium alone or in co-culture for 24h. This allowed us to determine how different growth environments affect expression of genes involved in nutrient acquisition and metabolism. Differentially expressed genes are provided in **Supplementary Files 1** (*F. nucleatum*) and **2** (*S. oris*). Principal components analysis (**Figs 2A** and **3A)** revealed that the greatest transcriptomic separation occurred between broth cultures media and airway epithelial co-cultures, with CFS exerting the most pronounced transcriptional shift among media conditions.

**Figure 2.**
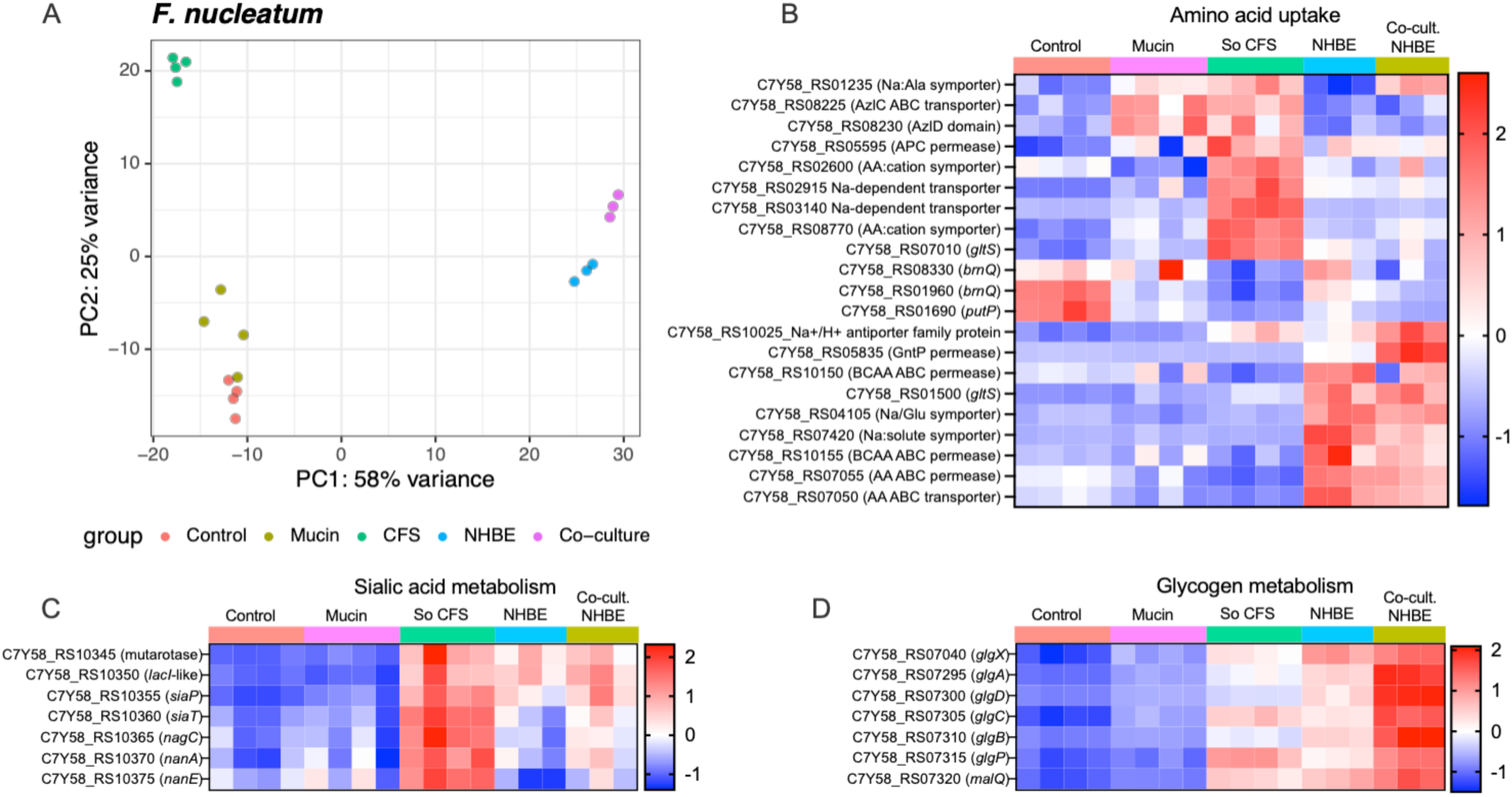
- Transcriptomic responses of *F. nucleatum* to mucin, *S. oris* supernatants, and epithelial co-culture. is. **A)** Principal component analysis (PCA) of *F. nucleatum* transcriptomes across control medium, mucin medium, CFS from *S. oris* grown in mucin medium, on normal human bronchial epithelia individually or in co-culture with *S. oris*. (**B)** Heatmap of amino acid transporter gene expression across conditions. (**C)** Expression of sialic acid catabolism (*nan* operon) genes. **(D)** Expression of glycogen metabolism genes (see Supplemental Files for complete gene lists). Each column in the heatmaps is an individual biological replicate and each row is a transcript. Normalized read counts from DESeq2 are scaled by row and presented as Z-scores.

**Figure 3.**
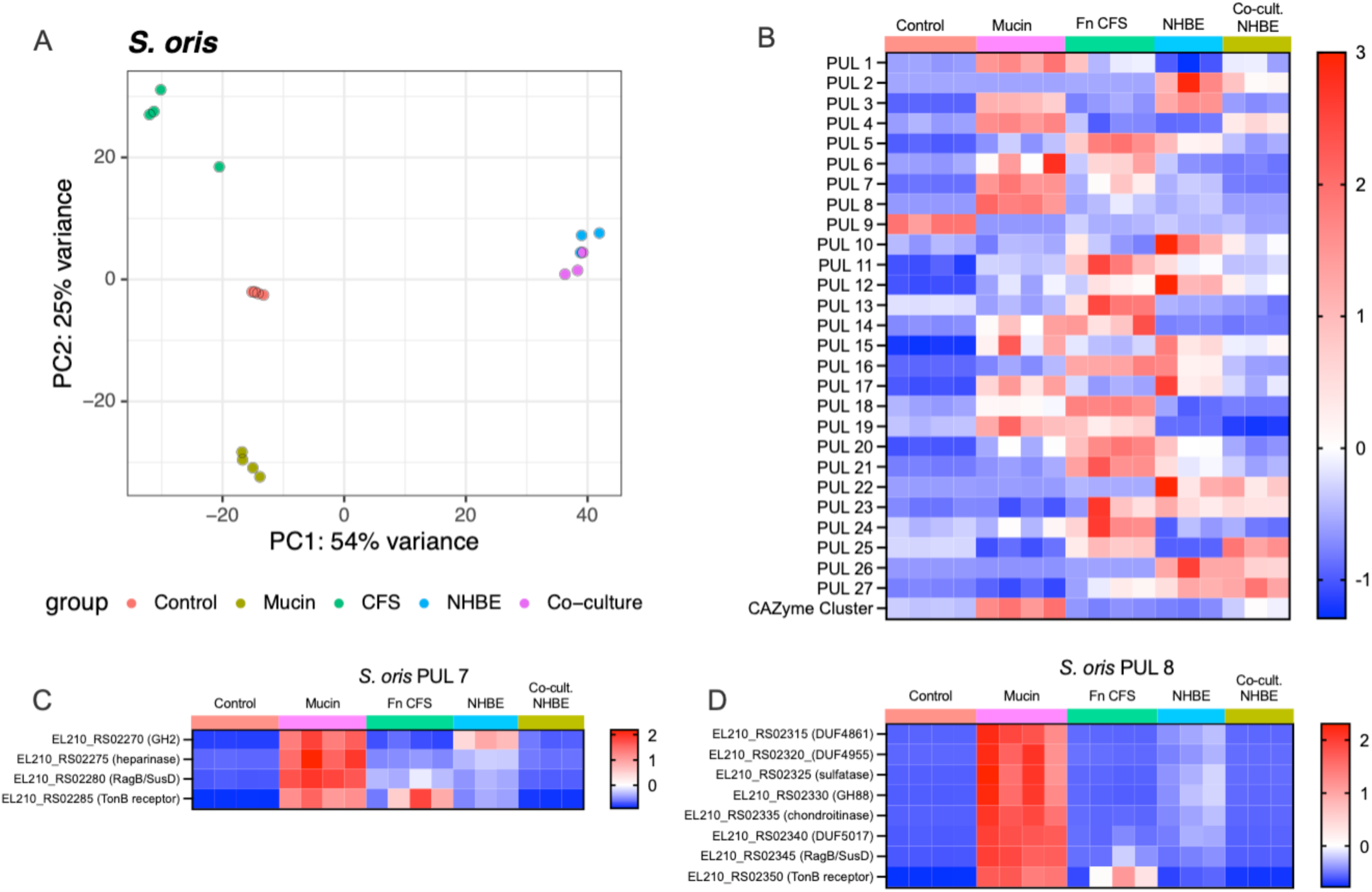
- The *S. oris* transcriptome is significantly altered across media and co-culture conditions *in vitro*. **A)** PCA of *S. oris* transcriptomes, colored by growth medium or co-culture status on NHBE. across growth conditions. lot depicting the transcriptome of individual *S. oris* samples, which are colored by growth medium or co-culture status on NHBE. Heatmaps of **B)** average expression of every gene in a given PUL across conditions, **C)** PUL 7, and **D)** PUL 8 in *S. oris*. Data are scaled by row and presented as Z-scores. Statistical significance for PUL expression comparisons provided in Suppl File Z.

### *F. nucleatum* amino acid uptake and metabolism

Since *F. nucleatum* preferentially ferments amino acids, we surveyed the expression of genes involved in amino acid uptake (**Fig. 2B**) and metabolism (**Suppl. Fig. 2**) to determine which substrates might be available across each growth medium or the airway epithelial environment. Two *brnQ* homologs encoding branched-chain amino acid (BCAA) importers and a *putP* proline transporter homolog were highly expressed in control medium, moderately expressed on NHBE, and showed the lowest expression in mucin, *S. oris* CFS (So CFS), and NHBE co-cultures. In contrast, homologs of the *azlC* and *azlD* BCAA importer family and a sodium:alanine symporter were upregulated in mucin and So CFS. Interestingly, the symporter exhibited low expression on NHBE alone but was elevated when *S. oris* was co-cultured on NHBE.

Several sodium-dependent symporters and transporters were specifically induced in So CFS, while multiple ABC-type transporters and permeases were selectively upregulated in NHBE co-culture, irrespective of *S. oris* presence. A sodium:proton antiporter and GntP-family permease were strongly induced in NHBE co-culture, but only in the presence of *S. oris,* suggesting interaction-dependent metabolic shifts. Genes involved in amino acid catabolism and short-chain fatty acid production were generally upregulated in NHBE co-culture with or without *S. oris*. This includes genes encoding butyrate production that have recently been linked to reactive oxygen/nitrogen resistance in *F. nucleatum* (40). Overall, while these transcriptional patterns suggest condition-specific substrate preferences, further experimental validation will be required to define precise substrate specificities.

### *F. nucleatum* carbohydrate metabolism is modulated by *S. oris*

Although *F. nucleatum* is known to catabolize glucose and fructose, other carbohydrate metabolic pathways were differentially expressed in So CFS and during co-culture with *S. oris* on NHBE cells. Since So CFS was derived from mucin-grown cultures, any glycan degradation by *S. oris* could supply the component sugars to *F. nucleatum*. For example, the *nan* operon (encoding a sialic acid catabolism pathway) was strongly induced in So CFS and in NHBE co-culture (**Fig. 2C**). Because *F. nucleatum* lacks a sialidase, its access to sialic acid appears dependent on exogenous glycosidase activity from *S. oris*. The absence of *nan* induction in mucin medium supports this model.

*F. nucleatum* also encodes genes annotated for glycogen production and degradation, though their function has not been experimentally confirmed. These genes were minimally expressed in control and mucin media but showed increased expression in So CFS and on NHBE, particularly when co-colonized with *S. oris* (**Fig. 2D**). More broadly, many genes predicted to encode carbohydrate active enzyme (CAZyme) genes showed additional differential expression across conditions (**Suppl. Fig. 3**), indicating that both host- and *S. oris-*mediated glycan degradation may shape *F. nucleatum* carbohydrate utilization, including genes involved in cell wall (e.g., *mrcB* homolog C7Y58_RS05695) and LPS assembly/modifications (e.g., *lpxCB* genes).

### Expression of *F. nucleatum* autotransporters and adhesins is modulated by nutrient and host conditions

Virulence in *F. nucleatum* is mediated by several Type V autotransporters that facilitate attachment to and invasion of host cells, and modulation of immune signaling (41–45). Therefore, we evaluated the expression of all annotated autotransporter genes across each broth and host-associated condition (**Suppl. Fig. 4**) Expression clustered into three groups: (i) genes whose expression is highest in the control medium, (ii) those with elevated expression in mucin or So CFS (e.g., *radD*), and (iii) genes with maximal expression on NHBE in both mono- and co-culture. The latter group included several well-characterized *F. nucleatum* virulence factor genes such as *fadA*, *cbpF*, fusolisin, and *fap2*. An interesting exception to this grouping was *fplA*, which was highly expressed in control, mucin, and NHBE co-culture, but was repressed in So CFS and in co-culture with *S. oris* on NHBE. These data are consistent with the increasing appreciation of the connections between bacterial metabolism and regulation of virulence factors and suggest that co-colonizing *S. oris* could modulate *fplA*-mediated contributions to *F. nucleatum* virulence (46).

### *S. oris* polysaccharide utilization loci (PUL) expression is responsive to both host and *F. nucleatum* interactions

The *S. oris* genome encodes 27 PULs and one putative CAZyme cluster, some homologous to validated systems in the *Bacteroides* genus (47,48). Such systems are used by bacteria to sense and degrade polysaccharides typically found in plant fibers (e.g., xylan) or host glycans like mucins (49). We averaged the expression of each gene in every PUL to determine which PUL(s) respond to glycans present in mucin and on the apical surface of airway epithelial cells, as well as whether *F. nucleatum* supernatants or co-culture on epithelia modulate PUL expression (**Fig. 3, Supplemental File 3 for ANOVA p-values**). All but PUL9 were minimally expressed in control medium. Eleven PULs and the CAZyme cluster were induced in mucin medium, consistent with mucin glycan utilization. Unexpectedly, PUL expression patterns were significantly altered in Fn CFS, with decreased expression of several mucin-induced PULs and induction of multiple mucin-insensitive PULs. A third group of PULs was induced during NHBE co-culture, including PULs 3, 5, 10, 12, and 15-17, many of which were suppressed in co-culture with *F. nucleatum*. In contrast, PULs 4 and 25, minimally expressed in NHBE monoculture, were induced by *F. nucleatum*. This variable expression of PULs was anticipated between growth in mucin and control media and on NHBE, but modulation of PUL expression in Fn CFS and co-culture on NHBE was unexpected. These data suggest that *F. nucleatum* may influence glycan foraging by *S. oris,* supporting a dynamic bidirectional interaction.

### *S. oris* encodes a putative Type VI secretion system

Like many members of the Bacteroidota, *S. oris* encodes a predicted contact-dependent multi-protein Type VI secretion system (T6SS) that delivers toxins to adjacent bacteria in intensely competitive conditions (50). Given that *S. oris* typically resides in polymicrobial settings and therefore could plausibly benefit from such a system, we scanned the *S. oris* NCTC 13071 genome for T6SS genes using SecReT6 (**Suppl. File 4**) (51). Three genomic loci were identified, including one containing 13 clear homologs of known T6SS genes, and two smaller clusters each containing four homologs. Expression profiles (**Suppl. Fig. 5**) revealed moderate expression in control medium and mucin media, with higher expression of select genes in Fn CFS. The *hcp* homolog was only expressed in control medium, while the *tssC* homolog was only expressed on NHBE, irrespective of the presence of *F. nucleatum*. A *tssI* homolog (locus tag EL210_RS03325) exhibited low-to-moderate expression across broth cultures and minimal expression in monoculture on NHBE but was highly induced in NHBE co-culture with *F. nucleatum,* suggesting potential functional activation in response to interspecies contact. Whether these genes constitute a functional T6SS will require further experimental validation.

### *F. nucleatum* drives airway epithelial inflammation, modulated by *S. oris* co-colonization

We next profiled the transcriptional response of NHBE cells colonized for 24h with *F. nucleatum* or *S. oris* singly or in co-culture, relative to uninfected controls (**Fig. 4A, Suppl. File 5**). PCA analysis revealed that *S. oris* alone had little effect on the host transcriptome, while *F. nucleatum*-induced a divergent host response. Co-colonization clustered separately from *F. nucleatum* colonization alone, suggesting *S. oris* modulates epithelial responses directly or indirectly via interactions with *F. nucleatum* that modulate its virulence. Gene set enrichment analysis (GSEA, **Fig. 4B, Suppl. Files 6 and 7**) identified enrichment of inflammation- and cancer-associated pathways in *F. nucleatum*-colonized epithelia, including *TNF-a*, IL-6/JAK/STAT3 signaling, IL-2/STAT5 signaling, and Wnt/β-catenin signaling. Co-colonization altered the magnitude and composition of these responses, highlighting the influence of microbial interactions on host inflammatory signaling. Indeed, co-culture of both species on NHBE also influenced expression of genes that were uniquely enriched relative to *F. nucleatum* mono-colonization. This is evident in **Fig. 4C,D**, where some inflammatory markers were induced by *F. nucleatum* and did not change in co-culture (e.g., *Il1β*), while others were only induced in the co-culture condition (e.g., *SQLE*, encoding squalene epoxidase). Together, these data reveal a complex interaction between all three species that is not apparent in NHBE cells mono-colonized with either species.

**Figure 4.**
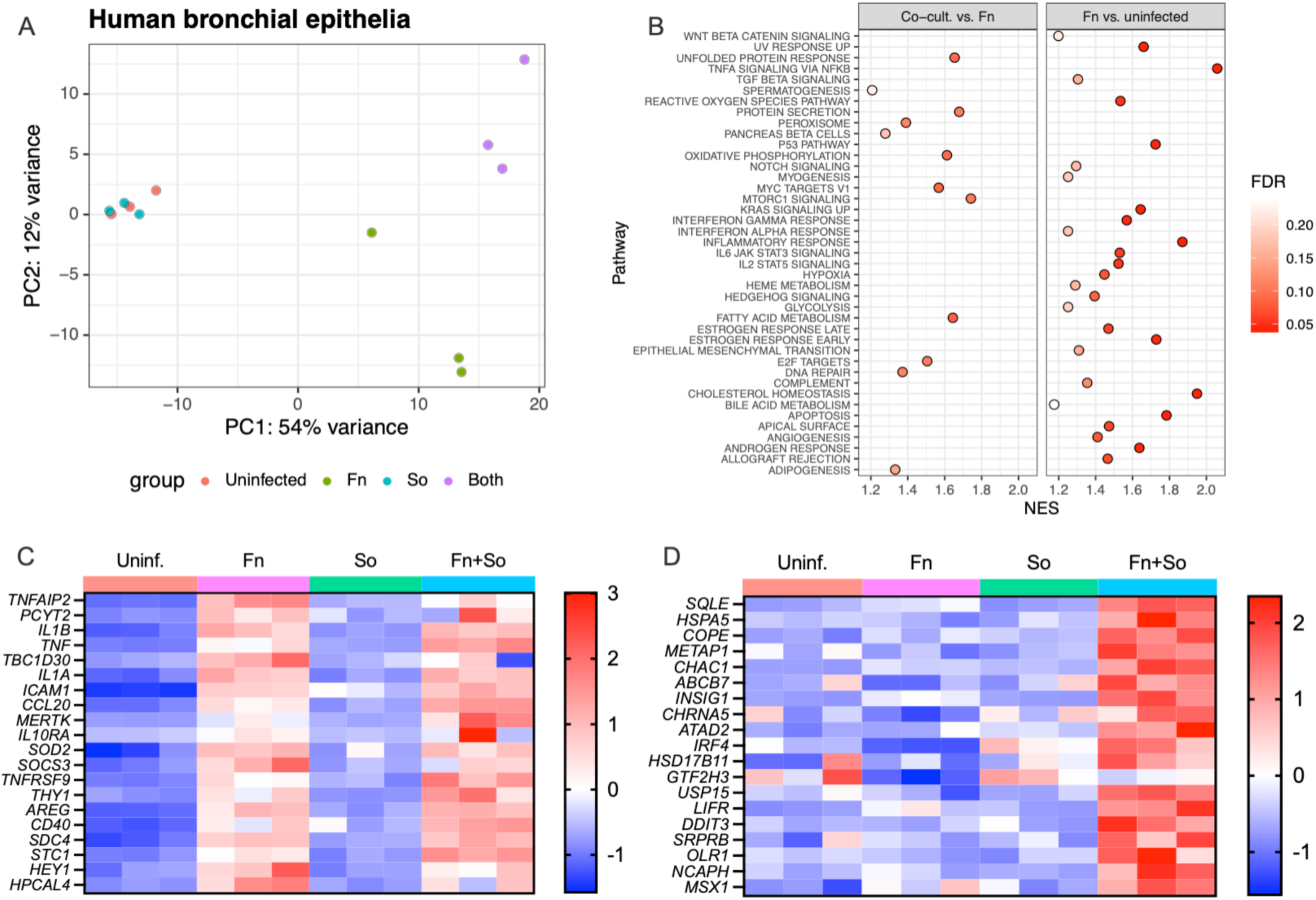
- Bacterial colonization alters the NHBE transcriptome. (**A)** PCA plot of host transcriptomes following mono- and co-colonization with *F. nucleatum* and *S. oris.* (**B)** Gene set enrichment analysis (GSEA) in samples from NHBE colonized with both *F. nucleatum* and *S. oris* compared to mono-colonization with *F. nucleatum* (left) and NHBE mono-colonized with *F. nucleatum* compared to uninfected cells. Normalized enrichment scores (NES) are shown on the x-axis and the color of each point indicates false discovery rate (FDR). High NES with low FDR indicates a pathway enriched in a given comparison. Pathways enriched in both comparisons were removed from the Co-cult vs. Fn plot to highlight pathways unique to this comparison, and only those pathways with a FDR < 0.25 are shown. Expression heatmaps of transcripts enriched in (**C)** cells mono-colonized with *F. nucleatum,* or (**D)** cells dual-colonized with *F. nucleatum* and *S. oris*.

## Discussion

The prevalence of *F. nucleatum* and *S. oris* co-colonization across multiple mucosal environments suggests potential for functional interactions that may impact bacterial colonization, persistence, and host responses. Using a combination of *in vitro* and host-relevant airway epithelial models, we show that these two microbes engage in bidirectional interactions involving glycan degradation, metabolite exchange, and transcriptional modulation. Although some overlaps were observed, the transcriptional profiles of both species in broth culture were markedly different from those observed during epithelial colonization. These differences were particularly evident in metabolic gene expression and the regulation of *F. nucleatum* virulence factors. These disparities underscore the importance of incorporating host-relevant conditions when using reductionist approaches to identify and model interactions between co-colonizing microbes.

Despite these differences, two transcriptional signals indicative of glycan-mediated interactions were observed across both culture and host-associated conditions. First, the *F. nucleatum nan* operon, encoding a sialic acid catabolism pathway, was not expressed in mucin medium but was highly induced in *S. oris* supernatants and in co-culture on NHBE cells. Given that *F. nucleatum* lacks sialidase activity, these data support a model in which sialic acid utilization depends on glycosidase activity from co-resident microbes. This is consistent with previous work by Agarwal et al (28), who showed that *F. nucleatum* exploits sialidase-positive microbes to access host-derived sialic acids in dysbiotic vaginal communities. The *nan* operon is not conserved across all *F. nucleatum* subspecies, so caution is warranted when generalizing these findings. Nevertheless, the broader concept of *F. nucleatum* contributing to dysbiotic bacterial community structures via cross-feeding is one that may apply to many sites of infection. Indeed, Queen et al identified *F. nucleatum* within polymicrobial biofilms on colorectal tumors and noted enrichment of sialic acid-adjacent metabolic pathways based on predicted metagenomic content (52).

A second glycan-associated transcriptional signal involved modulation of *S. oris* PULs. While several PULs were robustly induced by mucin, their expression was suppressed in the presence of *F. nucleatum* supernatants or during co-colonization on epithelial cells. This is notable for at least two reasons: (i) the large number of PULs affected; and (ii) the fact that *F. nucleatum* has not been reported to degrade host-associated glycans and is therefore not expected to alter the glycan landscape of mucin medium. The porcine gastric mucin used in this study is a crude proteolytic digest and likely contains non-mucin glycans such as glycosaminoglycans, which may explain the broad induction of PULs. For example, PULs predicted to target heparin and chondroitin (e.g., PULs 7 and 8) were induced in both mucin and on NHBE cells, but were repressed in the presence of *F. nucleatum,* suggesting nutrient depletion or transcriptional repression. One possible mechanism is that *F. nucleatum* secretes metabolic byproducts that modulate *S. oris* gene expression. This could include secreted polyglucose or glycogen, which could then be available to *S. oris* or scavenged from dead *F. nucleatum* cells. Notably, *F. nucleatum* glycogen metabolism genes were upregulated on NHBE, particularly during co-culture with *S. oris*, but not in mucin medium, suggesting that glycogen is unlikely to be present in *Fusobacterium* supernatants. Alternatively, glycosylated surface or secreted proteins from of *F. nucleatum* may serve as nutrient sources (53). A final possibility is that *F. nucleatum* encodes as-yet undescribed polysaccharide degradation and acquisition pathways that target glycans present in mucin or on epithelial cells, thereby diminishing PUL-inducing signals for *S. oris*.

The airway epithelial response was strongly influenced by *F. nucleatum* and further modulated by co-colonization with *S. oris.* Consistent with prior single-cell RNA-seq studies (39), *F. nucleatum* elicited robust inflammatory signaling, including induction of TNF-α, IL-1β, and mitochondrial stress responses suggestive of apoptosis. In contrast, *S. oris* alone triggered minimal host transcriptional changes, consistent with its commensal nature. Nevertheless, co-colonization with *S. oris* altered the epithelial response, suggesting that microbial interactions can modulate host sensing and inflammation. For example, *TNS4*, a gene linked to poor prognosis in lung adenocarcinoma (54), was significantly downregulated in the NHBE co-culture condition compared *F. nucleatum* alone. The opposite pattern is shown in **Fig. 4D**, where co-culture induced the expression of several genes not responsive to either bacterium alone. Because *S. oris* alone had minimal effect, we hypothesize that co-colonization alters *F. nucleatum* behavior in ways that modulate host responses. However, we cannot rule out *F. nucleatum*-driven changes in *S. oris* behavior that directly influences the airway epithelial response. Regardless, these data highlight the need to consider microbial interactions when modeling host-pathogen dynamics and support the integration of microbiome-derived hypotheses with mechanistic approaches to uncover host-pathogen-microbiome interactions.

In summary, we provide evidence that interactions between nutrient availability, the host environment, and co-colonizing bacteria shape the behavior of *F. nucleatum, S. oris,* and airway epithelial cells. Limitations should be acknowledged. While transcriptional and translational output are generally linked, differences between the half-life of mRNA and protein may result in temporal or spatial differences in bacterial and host behavior that are undetectable in our data. Likewise, site-directed mutagenesis is required to confirm several bacterial interactions suggested by our data, but genetic tools for *F. nucleatum* have only recently been developed and none are available for *S. oris* thus far. Lastly, we used only one strain of each bacterium in these experiments, which inherently excludes subspecies- and strain-level genetic (and thus phenotypic) diversity. Despite these limitations, our findings provide a framework for exploring glycan-driven cross-feeding and its impact on pathobiont behavior and host responses in mucosal environments.

## ACKNOWLEGEMENTS

This work was funded by the National Institute for Allergy and Infectious Diseases (AI177613), the University at Buffalo Research Foundation, and a Cystic Fibrosis Foundation postdoctoral fellowship to J.R.F. (002569F221)

