## Supplementary Data for "Cross-feeding interactions between *Fusobacterium nucleatum* and the glycan forager *Segatella oris*"

### Supplemental Data

**J.R. Fletcher and R.C. Hunter**

**Document S1.** Figures S1-S5

**S1 File.** *F. nucleatum* differentially expressed genes

**S2 File**. *S. oris* differentially expressed genes

**S3 File.** *S. oris* PUL ANOVA p-values

**S4 File.** *S. oris* SecReT6 predicted T6SS genes

**S5 File.** NHBE differentially expressed genes

**S6 File.** NHBE + Fn vs. NHBE untreated differentially expressed genes

**S7 File.** NHBE + Fn + So vs. NHBE untreated differentially expressed genes

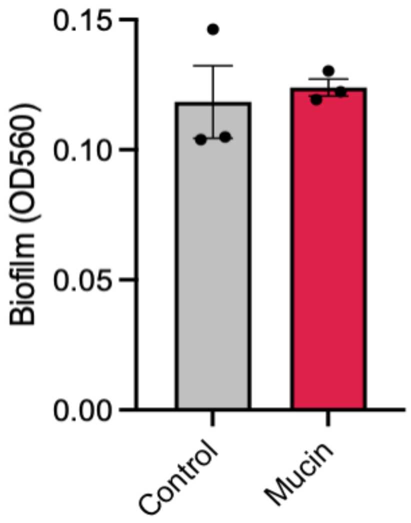


**Supplemental Figure 1 - Co-culture biofilms phenocopy *S. oris* mono-culture biofilms.** *F. nucleatum* and *S. oris* were grown together in control and mucin medium and biofilms were stained with crystal violet and quantified at OD560.

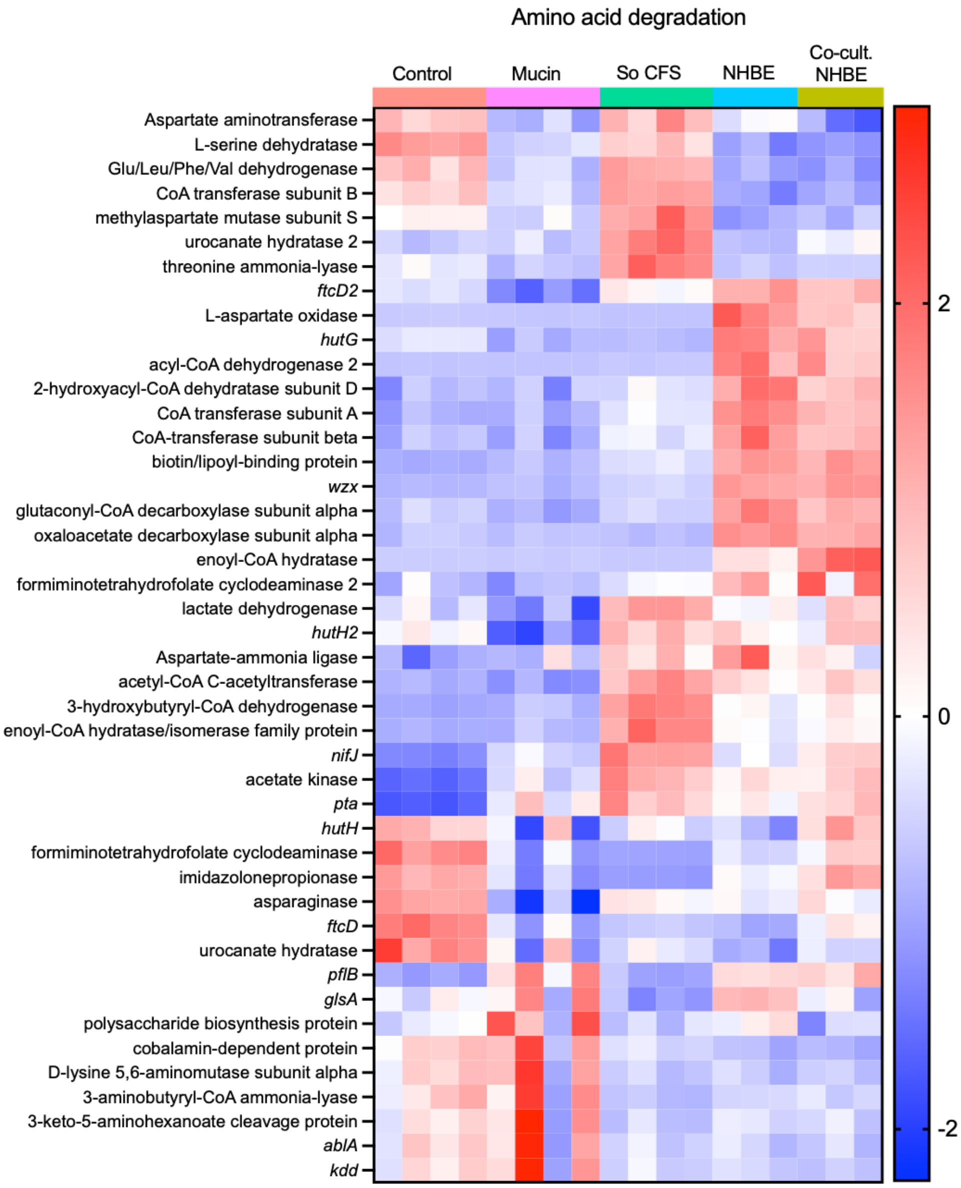


**Supplemental Figure 2** - Amino acid degradation gene expression across various media and on NHBE with and without co-culture with *S. oris*. Data are scaled by row and presented as Z-scores.

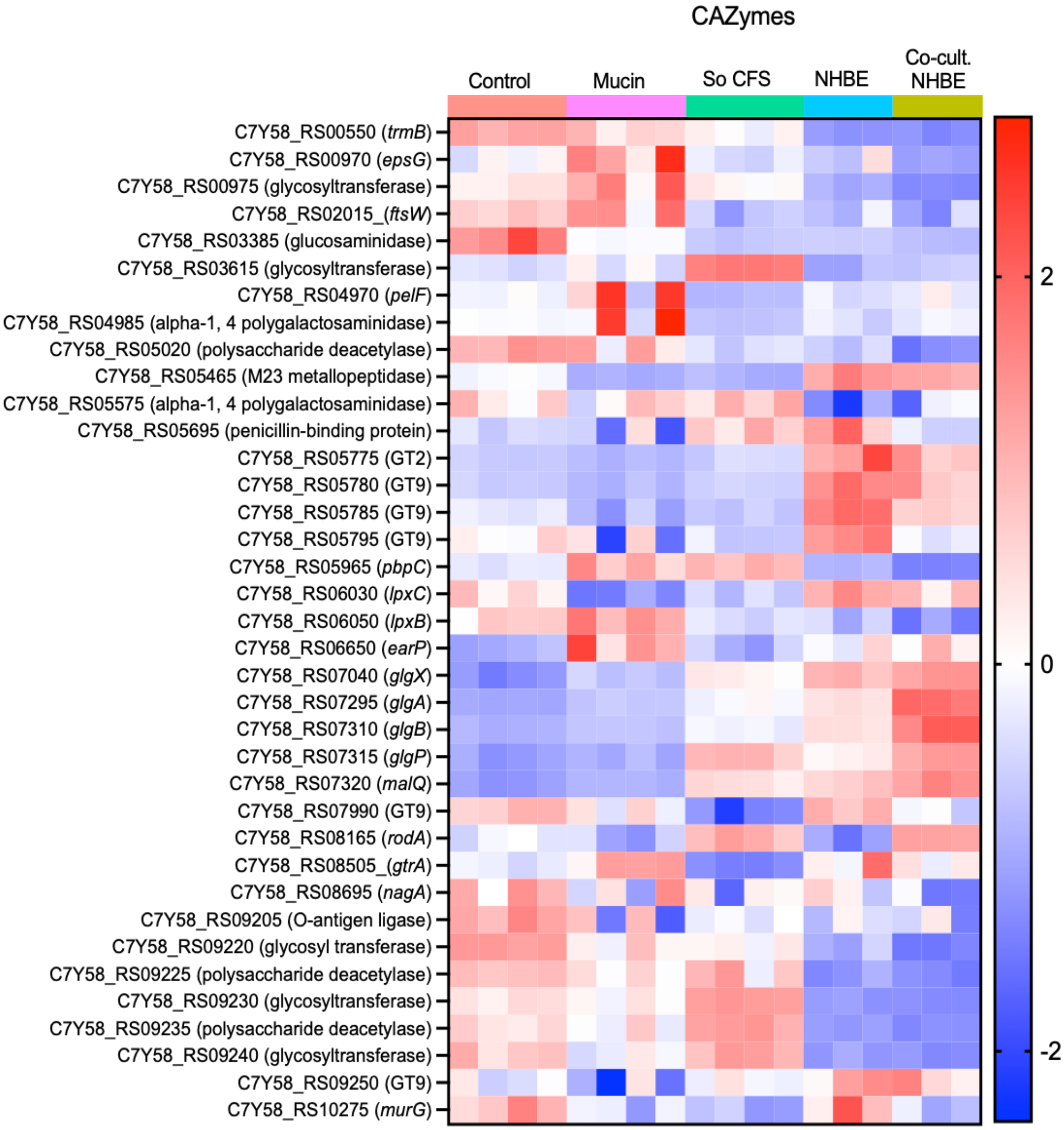


**Supplemental Figure 3** - Expression of *F. nucleatum* carbohydrate active enzymes (CAZymes) across various media and host conditions. Data are scaled by row and presented as Z-scores.

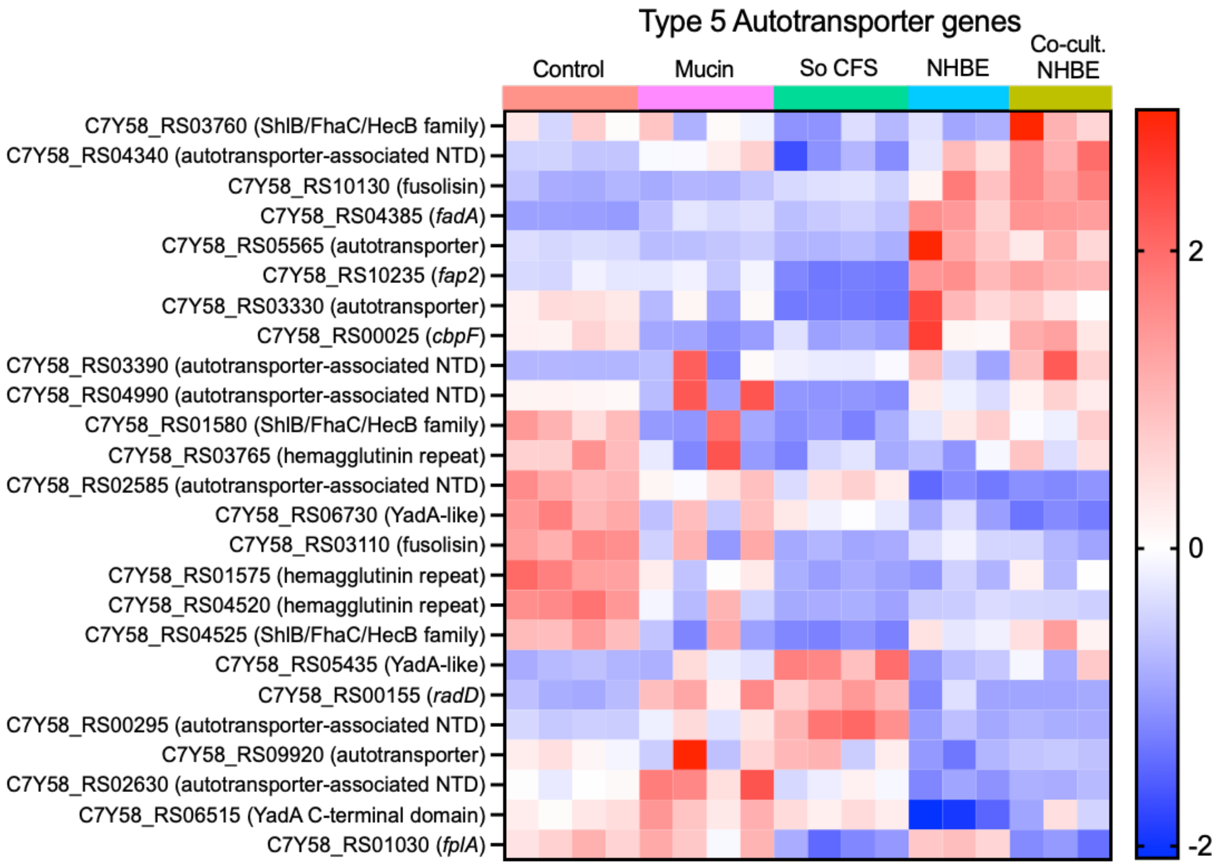


**Supplemental Figure 4 -** Expression of known and putative *F. nucleatum* adhesins and virulence factors. Data are scaled by row and presented as Z-scores.

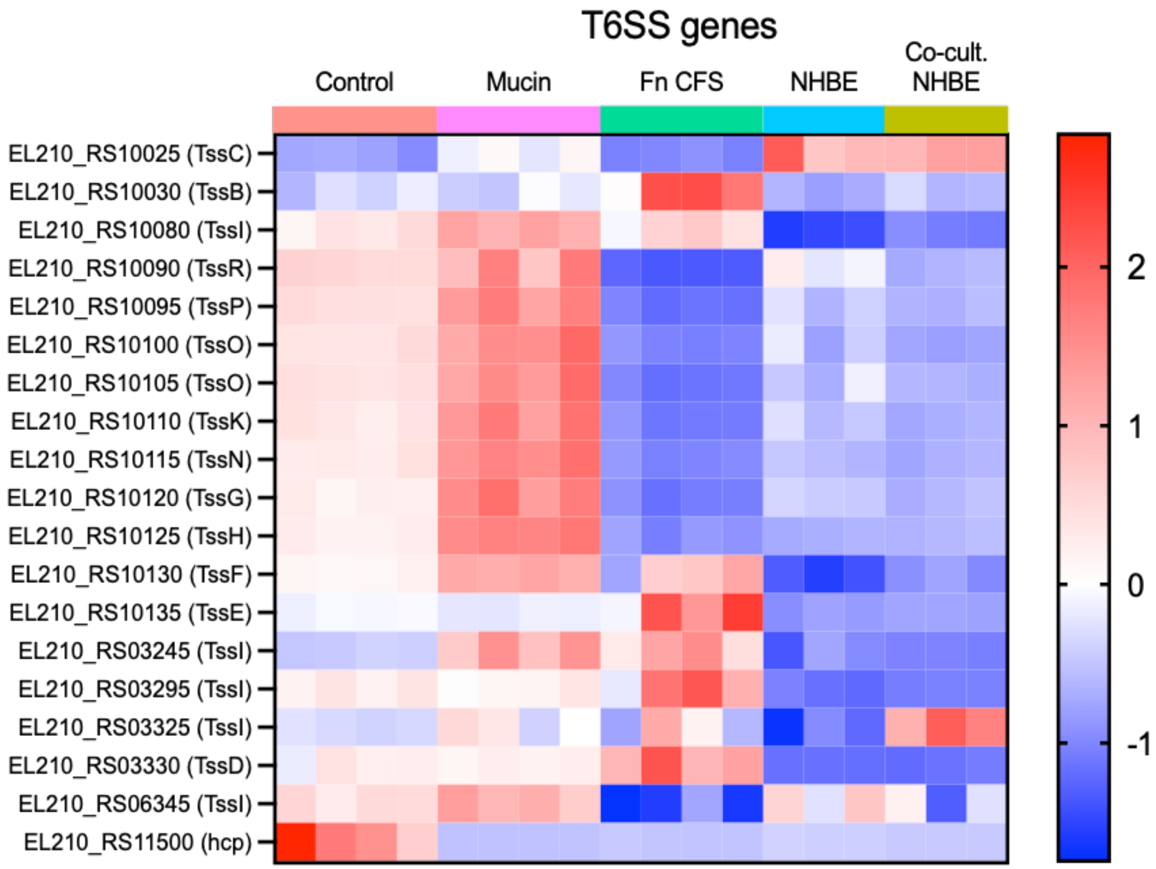


**Supplemental Figure 5 -** Expression of *S. oris* genes annotated as encoding Type VI Secretion System components across growth media and co-culture conditions on NHBE. Data are scaled by row and presented as Z-scores.
